# Kmo restricts *Salmonella* in a whole organism infection model by promoting macrophage lysosomal acidification through kainate receptor antagonism

**DOI:** 10.1101/2025.06.09.658567

**Authors:** Emily R. Goering, Anne E. Clatworthy, Margarita Parada-Kusz, Josephine Bagnall, Deborah T. Hung

## Abstract

The kynurenine pathway of tryptophan degradation has been implicated in various diseases, including cancer, neurodegenerative disorders, and infectious diseases. A key branchpoint in this pathway is production of the metabolite 3-hydroxy-kynurenine (3-HK) by the enzyme kynurenine 3-monooxygenase (Kmo). We have previously reported that administration of exogenous 3-HK promotes survival of zebrafish larvae to *Salmonella* Typhimurium infection by restricting bacterial expansion via a systemic mechanism that targets kainate sensitive glutamate receptor (KAR) ion channels and that the endogenous production of 3-HK by Kmo is required for defense against systemic *Salmonella* infection. Here we show that endogenous 3-HK promotes lysosomal acidification to contribute to macrophage microbicidal activity, with its absence leading to increased host susceptibility to infection. Further, 3-HK promotes lysosomal acidification in a KAR-dependent manner. We thus reveal a novel link between KARs and macrophage lysosomal acidification, and a novel mechanism by which 3-HK promotes control of bacterial infection.

**Author Summary:** Standard therapy for bacterial infections involves antibiotics to clear pathogens. However, the host immune system can also efficiently eliminate bacteria. We have recently shown that a metabolite of the kynurenine metabolic pathway, 3-hydroxy-kynurenine (3-HK), plays a role in the innate immune response to bacterial infection. Here, we show that the kynurenine pathway promotes macrophage clearance of intracellular bacteria by increasing lysosomal acidification of engulfed bacteria and that 3-HK does so by antagonizing kainate receptors. Together, this adds to our understanding of how multiple biological systems, including metabolic and immune pathways, interact to boost defense against bacteria.

## Introduction

Bacterial infection involves a complex dynamic between the pathogen and host that begins with host sensing of the bacterium followed by the triggering of a host response to ideally combat and clear the pathogen. The outcome of infection is thus determined by numerous mechanisms that define the balance between the host and pathogen. We had previously performed a whole organism chemical screen for small molecules that tip the balance in favor of host survival after infection not by acting directly on the pathogen but instead by modulating the host response. We discovered a small molecule, 3-hydroxy-kynurenine (3-HK), which upon exogenous administration to *Danio rerio* embryos infected with the gram negative pathogens *Pseudomonas aeruginosa* and *Salmonella* Typhimurium rescued the host from lethal systemic infection and decreased *in vivo* bacterial burden while having no effect on the bacteria in axenic culture (1). Exogenous 3-HK antagonizes kainate-sensitive ionotropic glutamate receptors (KARs), glutamate gated ion channels, to effect its host survival benefit (1). 3-HK’s control of bacterial burden within the entire organism could not be recapitulated in reductionist cell culture models, suggesting that a more complex integration of immune responses within the entire organism may be required for 3-HK’s protective effect (1).

Interestingly, 3-HK is also an endogenous metabolite produced by all eukaryotic cells in the kynurenine pathway, which metabolizes tryptophan to nicotinamide adenine dinucleotide (NAD^+^). The kynurenine pathway degrades more than 95% of dietary tryptophan and involves a number of enzymes beginning with indoleamine-2,3 dioxygenase (Ido1) or tryptophan 2,3-dioxygenase (Tdo), which converts L-tryptophan to N-formyl-kynurenine and then kynurenine (2) (Fig. 1A). Of these enzymes, Ido1 is induced in response to interferon-γ and other inflammatory stimuli, including inflammatory cytokines and TLR ligands, and is expressed in myeloid cells, endothelial cells and fibroblasts (3–6). Kynurenine is then hydroxylated by kynurenine 3-monooxygenase (Kmo) to form 3-HK (Fig. 1A), the metabolite we found which confers a survival benefit in lethal bacterial infection.

**Figure 1:**
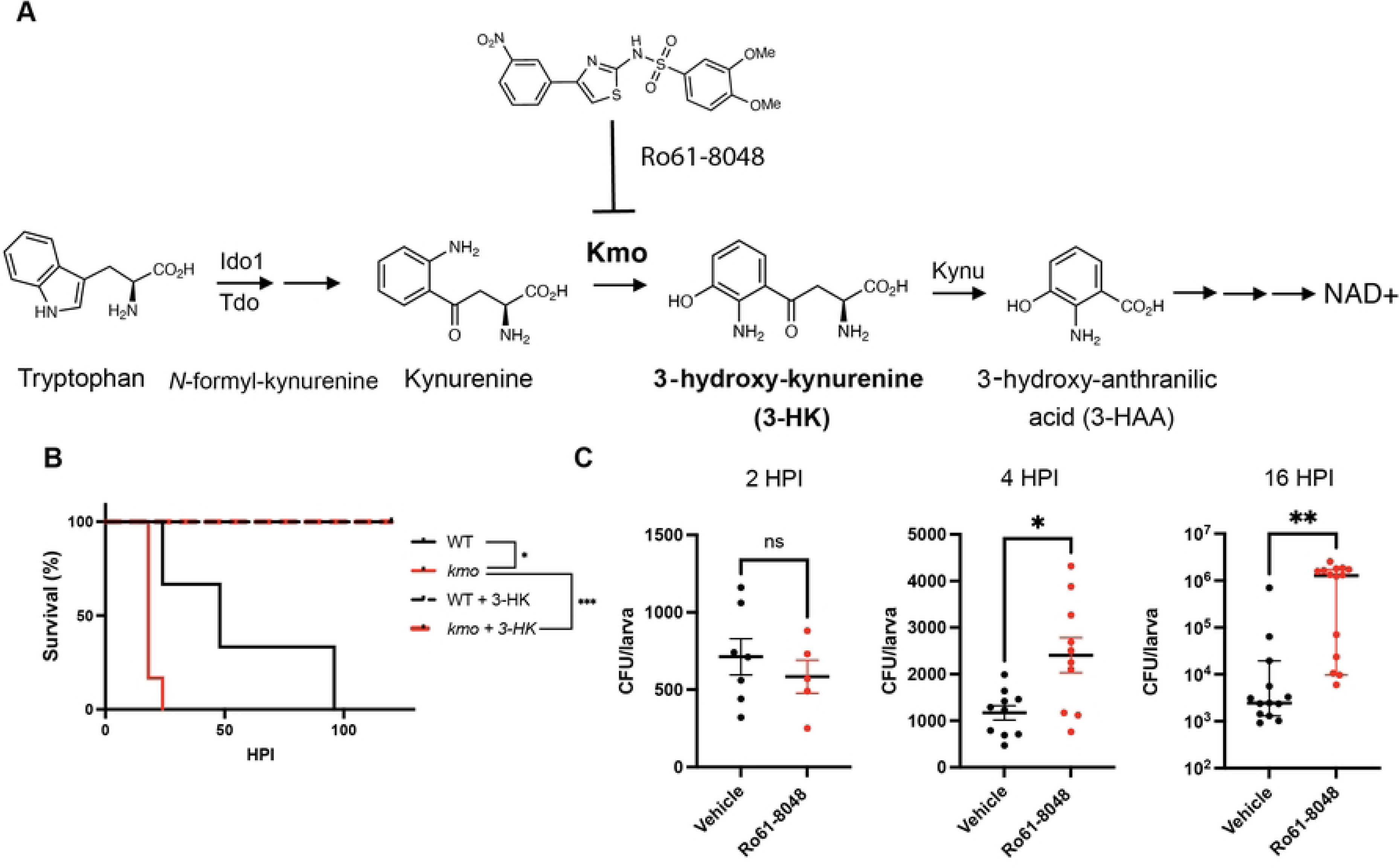
Endogenous production of 3-HK by Kmo is required for defense against ***S*. Typhimurium infection.** (**A**) The endogenous eukaryotic kynurenine pathway for tryptophan degradation that generates 3-HK (Ro61-8048 is a Kmo inhibitor). (**B**) Homozygous *kmo* mutant larvae infected 3 dpf with *S.* Typhimurium (CFU = 255 +/− 20) are sensitized to infection compared to their wildtype (WT) siblings. Sensitivity to infection is rescued by exogenous addition of 100µM 3-HK (N= 8 larvae/group; *p<0.05, ***p<0.001. Log rank test). (**C**) Larvae (3 dpf) infected with *S*. Typhimurium (CFU = 586 +/− 165) and treated with the Kmo inhibitor Ro61-8048 are impaired in their ability to control total organism bacterial burden as early as 4 hours post infection (HPI) compared to vehicle DMSO control. Data are representative of 3 experiments. CFU were quantified from individual larvae at indicated HPI; Unpaired t-test, *p<0.05, **p<0.01. dpf, days post fertilization; CFU, colony forming unit.

Inhibition of the endogenous kynurenine pathway and its production of 3-HK within zebrafish larvae increased susceptibility to infection. This hypersensitization could be reversed by the addition of exogenous 3-HK, thus implicating its endogenous production in defense against infection (1). The kynurenine pathway has been linked to host immunity against intracellular pathogens such as *Chlamydia trachomatis*, *Toxoplasma gondii, Listeria monocytogenes* and *Trypanosoma cruzi* through a number of mechanisms including aryl hydrocarbon receptor signaling, tryptophan depletion to control intracellular bacterial burden, or direct toxic effects of downstream metabolites (7–10). We identified this additional novel mechanism during lethal *S.* Typhimurium infection of zebrafish larvae as involving an interaction between endogenously produced 3-HK and KARs ion channels (1).

Here, we sought to understand how endogenously produced 3-HK plays a role in the innate immune defense against systemic *in vivo S*. Typhimurium infection. We used chemical or genetic inhibition of the enzyme Kmo as a means to block endogenous 3-HK production and examined the impact on macrophage control of infection, given that *S.* Typhimurium is known to utilize macrophages as a key intracellular niche for replication and systemic dissemination (11, 12). We found that knocking down *kmo* not only increases whole organism bacterial burden early in infection, but specifically the intracellular bacterial burden in individual macrophages, indicating a decrease in macrophage microbicidal activity. The impact of Kmo depletion on the expression programs of *S.* Typhimurium infected macrophages led to the hypothesis that inhibition of endogenous 3-HK could be impacting lysosome function. Utilizing microscopy and flow cytometry analyses, we confirmed that Kmo inhibition causes a defect in macrophage lysosomal acidification that was restored upon exogenous 3-HK application. Finally, we determined that macrophage lysosomal acidification was linked to KARs by demonstrating that KAR antagonism rescued the defect in lysosomal acidification caused by Kmo depletion. Together, this work sheds light on the mechanism by which the endogenous production of 3-HK promotes macrophage defense to bacterial infection.

## Results

### Endogenous production of 3-HK is required for control of bacterial burden in zebrafish larvae

We had previously shown that knockdown of *kmo* with a translation blocking morpholino or chemical inhibition of Kmo with the known inhibitor Ro61-8048 sensitized animals to infection compared to control or vehicle treated animals, in a manner that could be rescued by chemical complementation of Kmo activity with the addition of exogenous 3-HK (Fig. S1B, S1C) (1). Here we further confirmed this phenotype by examining this phenotype in CRISPR/Cas9 *kmo* knock-out larvae (Fig. 1A). We created an allele of *kmo* that contains an 82 base pair deletion in exon 7 using CRISPR/Cas9, resulting in a truncated protein and significantly reduced *kmo* expression in homozygous mutant animals relative to wildtype by RT-qPCR (Fig. S1A). Infection of offspring from heterozygous parents at 3 days post-fertilization (dpf) revealed that homozygous *kmo* mutant offspring are significantly sensitized to sublethal intravenous challenge with *S.* Typhimurium compared to their wildtype siblings (Fig. 1B).

Importantly, chemical genetic complementation of Kmo activity with exogenous 3-HK rescued the sensitivity to infection observed following *kmo* knock-out (Fig. 1B, Fig. S1B, and Fig. S1C). Together, the three orthologous methods, *kmo* morphant, chemical Kmo inhibition, and now CRISPR/Cas9 knockout, independently confirmed that endogenous production of 3-HK by Kmo is required for survival to systemic bacterial infection with *S*. Typhimurium in zebrafish.

To elucidate how endogenous production of 3-HK promotes survival following systemic *S.* Typhimurium infection, we first enumerated bacteria from Kmo inhibitor and vehicle treated larvae over time to determine the impact of Kmo inhibition on total bacterial burden. Kmo inhibitor treated larvae displayed a more rapid expansion in bacterial burden as early as 4 hours post infection (HPI) and by 16 HPI had over a 100-fold increase in bacterial load per animal compared to vehicle treated fish (Fig. 1C). While 3-HK is known to have pro-oxidant activity, the inability of Kmo inhibitor treated animals to control bacterial burden could not be attributed to a defect in ROS generation, as treatment with the Kmo inhibitor resulted in enhanced ROS production in the fish over time compared to vehicle control (Fig. S2A). Likewise, Kmo inhibitor treatment did not impair production of pro-inflammatory cytokines Tnf-α and Il-1β, which were also increased at late stages of infection in treated fish (Fig. S2B, Fig. S2C). Thus, the decreased survival and loss of control of bacterial burden when the production of endogenous 3-HK is inhibited is not accounted for by impacts on ROS and pro-inflammatory cytokine levels.

### Endogenous 3-HK promotes control of bacterial burden within macrophages

At three days post infection, the larval immune system consists of an active innate immune system that is primarily composed of macrophages and neutrophils, while adaptive immunity is not yet fully developed (13). Multiple studies have shown that macrophages are critical to the early bloodstream response to *S.* Typhimurium infection in fish (14–16). The majority of S. Typhiumurium bacteria are located within macrophages within the first 4 hours of infection, where bacteria can either be eliminated or allowed to replicate depending on macrophage function (17). We also showed that macrophages are required for exogenous 3-HK’s effect on larval survival (1). We thus wondered if inhibition of endogenous 3-HK production could be impairing the macrophage response to infection.

In principle, poor control of bacterial burden when endogenous production of 3-HK is impaired could result from 1) a decrease in macrophage numbers in zebrafish larvae, as this has been shown to increase sensitivity to infection (1, 18); 2) decreased phagocytic capacity of macrophages; or 3) decreased microbicidal activity of macrophages. To examine each of these possibilities, we first quantified macrophage numbers by microscopy using a Tg(*mpeg1.1:mCherry*) zebrafish line where macrophages are fluorescently labeled with mCherry (19). Comparing *kmo* morphant and control larave, and Kmo inhibitor and vehicle treated larvae, we found no differences in macrophage numbers between Kmo inhibited or knockdown larvae compared to controls in both uninfected and infected animals (Fig. 2A, Fig. S3A, Fig. S3B, Fig. S3C). Next, we evaluated the phagocytic capacity of macrophages by injecting larvae with GFP+ *S*. Typhimurium or GFP+ latex beads. We found no differences in the percentage of macrophages which had taken up GFP+ bacteria or GFP+ latex beads by flow cytometry at 4 hours post-infection following Kmo inhibition or in *kmo* knockdown larvae versus control (Fig 2B, Fig. S3D, Fig. S3E, Fig. S3F). Thus, decreased macrophage numbers or impaired macrophage phagocytosis does not account for increased bacterial load resulting from inhibition of endogenous 3-HK production.

**Figure 2:**
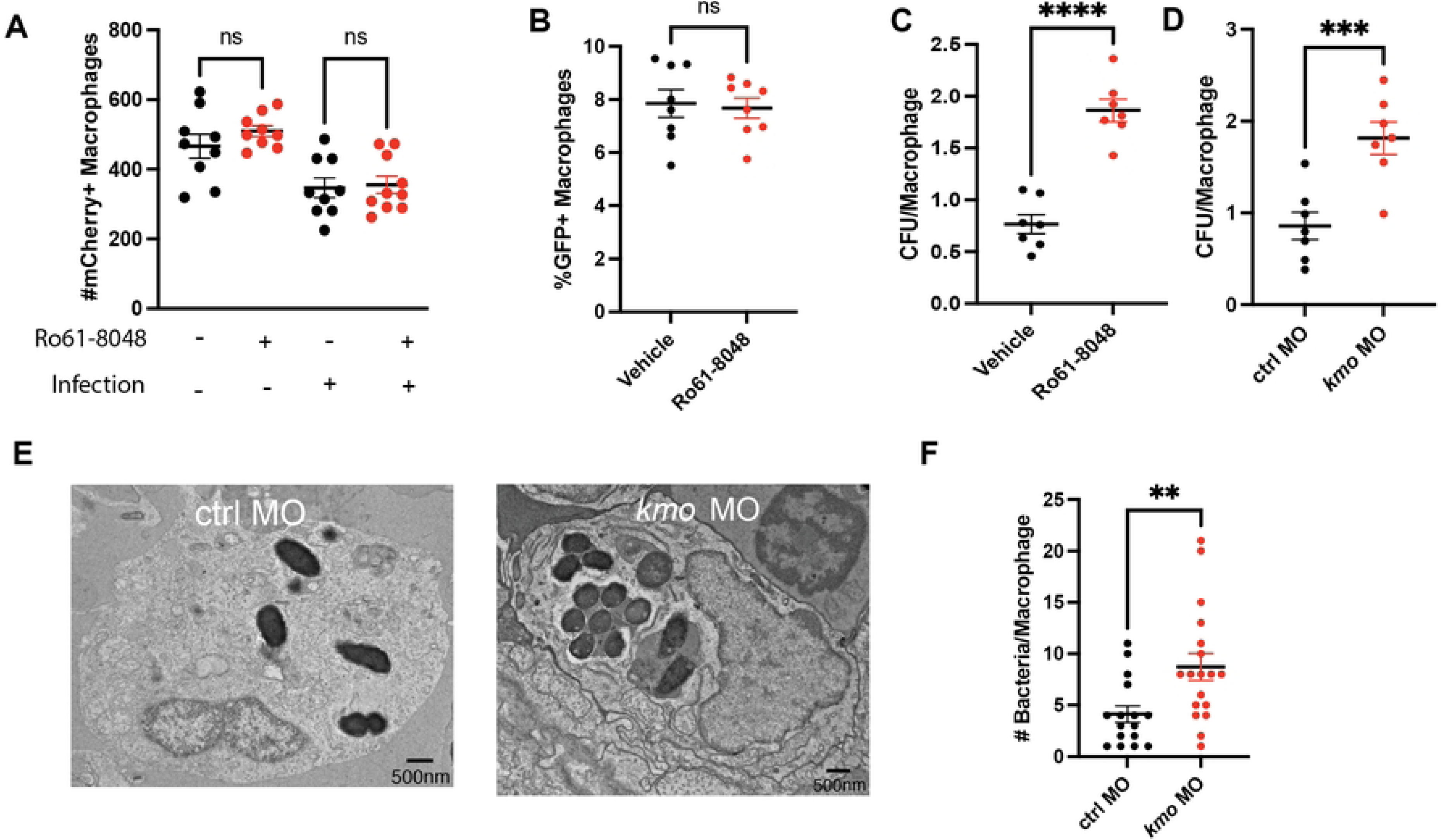
K**m**o **is required for intracellular control of bacterial burden within macrophages.** (**A**) Treatment with the Kmo inhibitor Ro61-8048 does not change macrophage numbers in either infected or uninfected larvae compared to DMSO control. Macrophage numbers from Tg(*mpeg1.1:mCherry*) larvae either treated with 50µM Ro61-8048 or DMSO control were quantified by microscopy from both infected (16 HPI) and sibling uninfected larvae (values, mean +/− SEM). Each data point represents an individual larva. Data are representative of 3 experiments. (One-way ANOVA followed by Tukey’s multiple comparisons, ns, non-significant). (**B**) Kmo inhibition with Ro61-8048 does not impair macrophage engulfment of *S*. Typhimurium-GFP. Tg(*mpeg1.1:mCherry*) larvae (3 dpf) were infected with *S*. Typhimurium-GFP and the percentage of mCherry+/GFP+ macrophages were quantified by flow cytometry at 4 HPI (Unpaired t-test, ns, non-significant). Each data point represents an individual larva. Data are representative of 3 experiments. (**C**-**F**) Kmo inhibition (**C**) or depletion (**D-F**) impairs macrophage control of bacterial burden. (**C**) Tg(*mpeg1.1:mCherry*) larvae (3 dpf) were infected with *S.* Typhimurium-GFP (CFU/ larvae) and treated with 50µM Ro61-8048 or DMSO control or (**D**) *kmo* or control morphant larvae were infected with *S.* Typhimurium-GFP (CFU/ larvae) and the number of live bacteria per sorted macrophage (mCherry+/ GFP+) was quantified by isolating macrophages by FACS at 4 HPI followed by lysis and enumeration of bacteria by plating for CFU (Unpaired t-test; C, ****p<0.0001, D, ***p<0.001). Data are representative of 3 experiments. (**E**, **F**) Kmo depletion increases numbers of intracellular bacteria per macrophage by transmission electron microscopy (TEM). *kmo* or control morphant larvae were infected with *S.* Typhimurium, fixed at 4 HPI and infected phagocytes in the caudal hematopoietic tissue were examined by TEM. (**E**) Representative transmission electron microscopy images. (**F**) Quantification of observed bacteria per phagocyte (Values, mean +/− SEM; Unpaired t-test, **p<0.01). Infected phagocytes were imaged and quantified from 3 biological replicates. Infected phagocytes were identified by the presence of black, electron dense bacteria. Each data point represents a single phagocyte. CFU, colony forming units. HPI, hours post infection. dpf, days post fertilization. Ro61-8048, Kmo inhibitor.

Finally, we examined whether inhibition of endogenous 3-HK production might impair the microbicidal capacity of macrophages. To examine this possibility, we quantified the number of live bacteria per macrophage in *kmo* knockdown and inhibitor treated larvae compared to controls utilizing fluorescence activated cell sorting (FACS) to isolate infected cells and then plating for colony forming units (CFU) to enumerate live bacteria. We infected Tg(*mpeg1.1:mCherry*) larvae with *S*. Typhimurium-GFP and sorted and lysed infected, double-positive mCherry+/GFP+ macrophages for intracellular bacterial burden (Fig. S4). We found that Kmo inhibition (Fig. 2C) or *kmo* knockdown (Fig. 2D) resulted in a significantly higher number of live *S*. Typhimurium per macrophage compared to controls. This finding is consistent with our previous results where exogenous 3-HK application had the opposite effect and resulted in fewer bacteria per macrophage compared to controls (1).

We confirmed this finding using transmission electron microscopy (TEM) of the caudal hematopoietic tissue, which contains a high proportion of larval macrophages at this developmental stage (20). Here we also found that *kmo* morphant animals displayed a higher number of bacteria per phagocyte compared with control morphant larvae (Fig. 2E, Fig. 2F). In the TEM images, we observed both double– and single-membrane vacuoles surrounding intracellular *S.* Typhimurium in phagocytes from both *kmo* morphant and control morphant larvae (Fig. S5). This observation differs from previous work suggesting that *S.* Typhimurium bacteria are primarily enclosed within single-membrane vacuoles in embryos infected 48 hpf (18) which might reflect differences in macrophage functionality between 72 hpf and 48hpf developmental stages.

Taken together, these results show that inhibition of endogenous production of 3-HK impairs the microbicidal capacity of macrophages after phagocytosis, resulting in an increase in bacterial burden per macrophage. Application of exogenous 3-HK has the opposite effect and promotes control of bacterial burden within macrophages (1).

### Endogenous production of 3-HK promotes lysosomal acidification in macrophages

To investigate why the loss of endogenous 3-HK results in increased bacterial burden in macrophages, we examined the transcriptional program of infected macrophages when endogenous 3-HK production is inhibited. We performed bulk RNA sequencing of infected macrophages to compare transcriptional changes with and without *kmo* depletion. We infected *kmo* and control morphants from Tg(*mpeg1.1:mCherry*) larvae with *S*. Typhimurium-GFP and isolated by FACS mCherry+/GFP+ double-stained infected macrophages at 4 HPI; we also isolated mCherry+ macrophages from uninfected, control larvae in parallel (Fig. 3A). We then performed transcriptional analysis on isolated populations. Gene set enrichment analysis (GSEA) confirmed, as expected, that functional pathways of the inflammatory response, defense response, response to cytokines, and regulation of immune system process were enriched in *S*. Typhimurium infected macrophages compared to uninfected macrophages (Fig. S6A, Fig. S6B)(1, 21). In addition, GSEA analysis using KEGG terms also found that the lysosomal pathway was significantly downregulated in infected *kmo* depleted macrophages compared to infected control macrophages (Fig. 3B, Fig. 3C), including genes encoding the vacuolar ATPase, lysosomal membrane proteins, and acid hydrolases. Hence, *kmo* knockdown results in a global suppression of genes necessary for formation and proper acidification of lysosomes, which could explain why the loss of endogenous 3-HK results in increased bacterial burden within macrophages.

**Figure 3:**
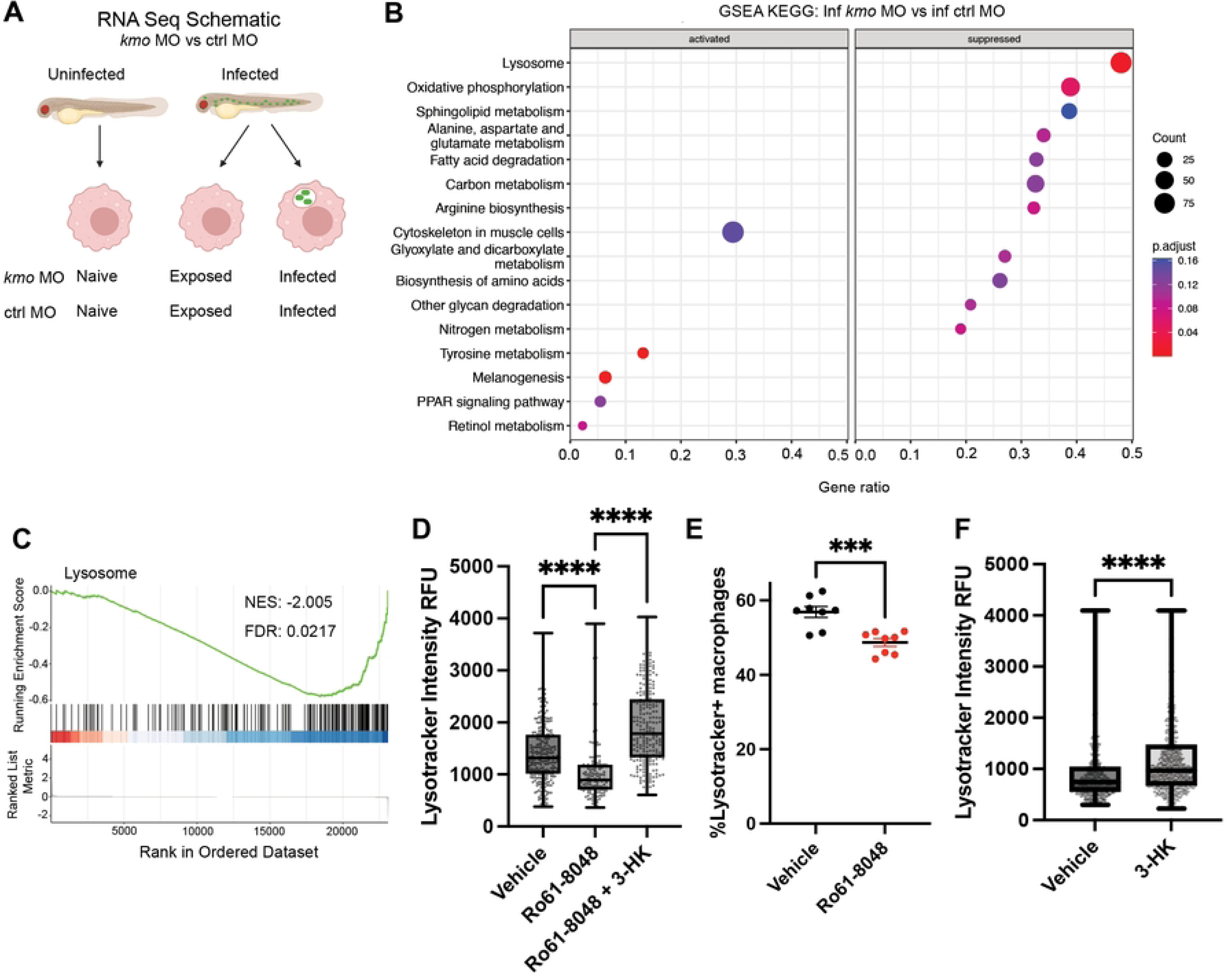
K**m**o **inhibition suppresses lysosomal acidification in macrophages during infection which is rescued by 3-HK.** (**A**) Schematic of bulk RNA sequencing measuring transcriptional changes in naïve, exposed and infected macrophages from infected *kmo* and control morphant larvae. Tg(*mpeg1.1:mCherry*) *kmo* or control morphant larvae were infected with *S*. Typhimurium-GFP or left uninfected and mCherry+ macrophages were sorted from infected and uninfected (naïve) fish at 4 HPI. Macrophages from infected fish were further sorted into infected (GFP+) and exposed (GFP-) populations. (**B-C**) *kmo* depletion suppresses expression of gene signatures of lysosomal acidification in infected macrophages compared to controls using GSEA (**B**, KEGG terms). **C**) GSEA plot showing downregulation of the lysosomal KEGG pathway in infected *kmo* macrophages compared to infected control macrophages. Enrichment score shows suppression of the lysosomal pathway, with lysosomal pathway genes predominantly clustered at the bottom of the ranked gene list. NES, normalized enrichment score. FDR, false discovery rate. (**D, E, F**) Endogenous and exogenous 3-HK promote lysosomal acidification in infected macrophages. Tg(*mpeg1.1:mCherry*) larvae were infected with *S.* Typhimurium-GFP and treated with DMSO control or 50µM Ro61-8048 with or without 100µM 3-HK. Embryos were bathed in Lysotracker to stain acidic compartments at 4 HPI and maximum Lysotracker fluorescence intensity was measured using confocal microscopy. Lysotracker intensity was quantified from imaged individual macrophages in the caudal hematopoietic tissue. Data are representative of 3 experiments. (**D**,**E**; Values are individual macrophages from 8-13 biological replicates; All data points shown; Unpaired t-test. ****p<0.0001). (**F**) Quantification of the percentage of total bright Lysotracker+ macrophages in whole larvae via flow cytometry in 50µM Ro61-8048 and DMSO (vehicle) control treated fish. (Each value represents pooled 5 larvae; Unpaired t-test, ***p<0.001). Data are representative of 2 experiments. GSEA, gene set enrichment analysis. ctrl, control morphant. *kmo*, *kmo* morphant. Ro61-8048, kmo inhibitor.

To functionally validate that loss of endogenous 3-HK impairs lysosomal function, we used the pH-sensitive fluorescent dye, Lysotracker, to measure lysosomal acidification in macrophages during infection. We infected Tg(*mpeg1.1:mCherry*) larvae with *S*. Typhimurium-GFP, stained animals with Lysotracker at 4 HPI, and measured lysotracker staining in mCherry+ cells using confocal microscopy and flow cytometry. We found a significant reduction in Lysotracker staining in both infected (mCherry+/GFP+) and uninfected (mCherry+/GFP-) macrophages from larvae treated with the Kmo inhibitor compared to vehicle control (Fig. 3D, Fig. 3E, Fig. S7A), demonstrating a global reduction in lysosome acidification regardless of infection status. Importantly, application of exogenous 3-HK rescued this defect in Lysotracker staining following Kmo inhibition and loss of endogenous production of 3-HK (Fig. 3D, Fig. S7A). Further, exogenous application of 3-HK also increased the maximum Lysotracker staining in wildtype macrophages compared to vehicle controls (Fig. 3F), consistent with an enhanced ability to control bacterial burden in macrophages in 3-HK treated, wildtype animals (1).

To further corroborate this data, we assessed the acidification capacity of macrophages using pHrodo-conjugated *Escherichia coli* which fluoresces brightly in low pH environments, indicative of *E. coli* localization within lysosomes. We injected heat-killed pHrodo-labeled *E.coli* into the bloodstream of Tg(*mpeg1.1:mCherry*) larvae and assessed the percentage of macrophages with bright pHrodo staining by flow cytometry 4 HPI. We found that Kmo inhibitor-treated fish (Fig. S7B) or *kmo* morphants (Fig. S7C) had a significantly lower percentage of macrophages staining brightly for pHrodo compared to control. Together, these data show that endogenous and exogenous 3-HK act to promote lysosomal acidification in macrophages.

Finally, we determined if lysosomal acidification was important for the survival effect of 3-HK. We treated infected and control larvae with the vacuolar ATPase (vATPase) inhibitor bafilomycin A1 (22). Bafilomycin A1 treatment not only sensitized larvae to infection, as expected, but it also completely abrogated the protective effect of 3-HK, indicating that lysosomal acidification is necessary for 3-HK to protect against infection (Fig S8). 3-HK thus mediates clearance of intracellular bacteria by affecting lysosome function.

### Endogenous production of 3-HK promotes lysosomal acidification via KARs

We had previously found that exogenous 3-HK acts through KARs to promote survival following systemic infection, with the known KARs antagonist NS 3763 also affording protection from lethal infection (1). KARs are a subclass of ionotropic glutamate receptors. They are ligand-gated ion channels activated by glutamate that are most commonly studied in the context of synaptic transmission in neurons (23). Here we confirmed that endogenous 3-HK also works via KARs since the KAR-specific antagonist NS 3763 reverses the sensitivity to infection afforded by *kmo* knockdown (Fig. 4A).

**Figure 4:**
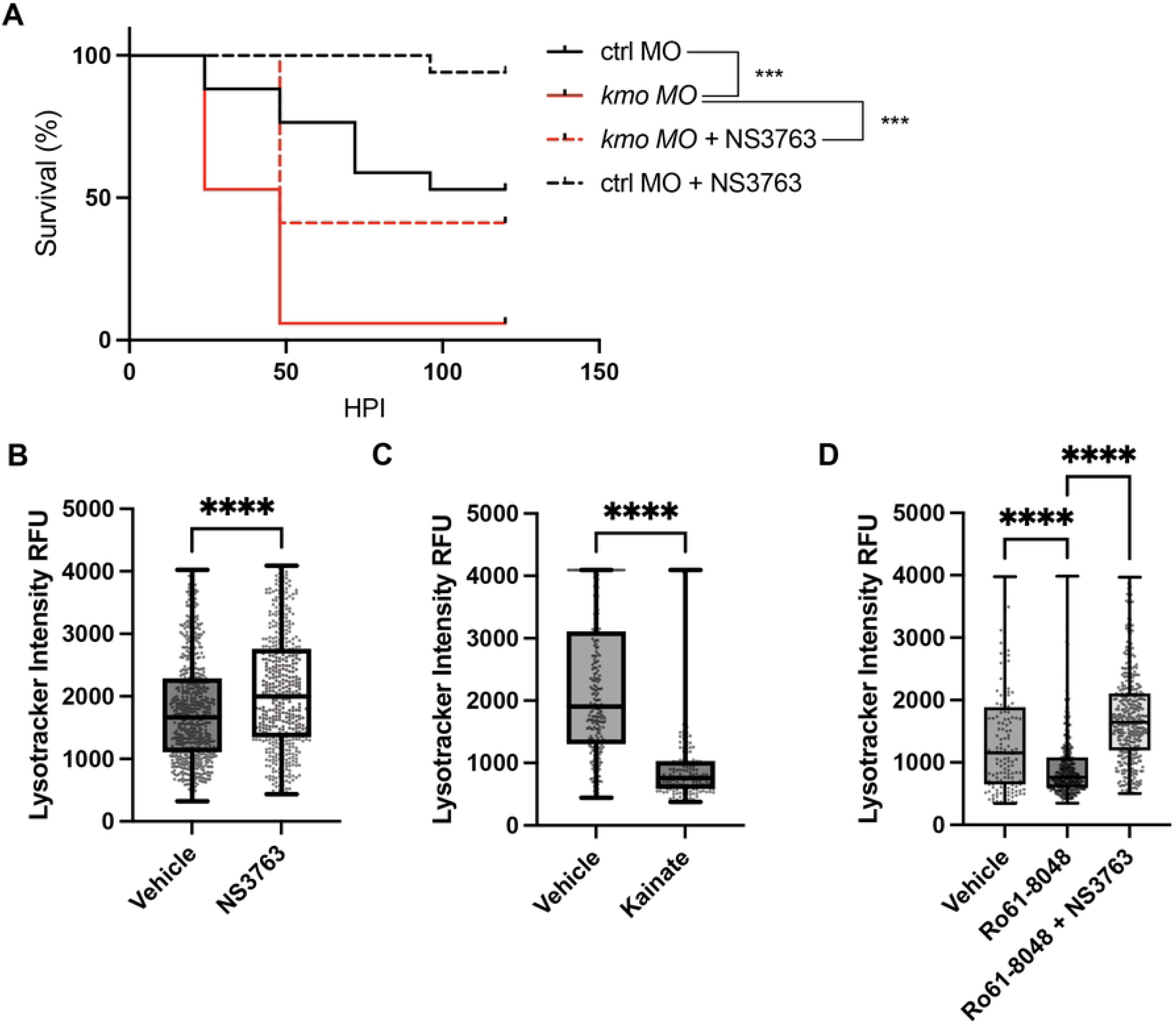
K**a**inate **receptor (KAR) antagonism rescues the loss of endogenous 3-HK and promotes lysosomal acidification in infected macrophages.** (**A**) KAR antagonism with NS 3763 rescues the sensitivity to infection in *kmo* morphants. Kmo and control morphant larvae were infected with *S.* Typhimurium (215 +/− 134 CFU), treated with 75µM NS 3763 or DMSO control and monitored over time for survival. (N= 17 larvae/ group; ***p<0.001, Log rank test). (**B, C, D**) Tg(*mpeg1.1:mCherry*) larvae were infected with *S.* Typhimurium-GFP and maximum Lysotracker staining was measured at 14 HPI using confocal microscopy. Lysotracker intensity was quantified from individual macrophages in the caudal hematopoietic tissue. (**B**) The KAR antagonist NS 3763 increases maximum Lysotracker staining in infected macrophage populations compared to vehicle (DMSO) control (Unpaired t-test. ****p<0.0001). (**C**) The KAR agonist kainate decreases maximum Lysotracker staining in infected macrophages compared to vehicle (DMSO) control (Unpaired t-test. ****p<0.0001). Values are individual macrophages imaged in the caudal hematopoietic tissue. All data points shown. (**D**) KAR antagonism with NS 3763 reverses the decrease in Lysotracker staining in macrophages following inhibition of endogenous 3-HK production with the Kmo inhibitor Ro61-8048. (One-way ANOVA followed by Tukey’s multiple comparisons. ****p<0.0001). Values are individual macrophages from 8-14 biological replicates. All data points shown.

Given this relationship between endogenous 3-HK and KARs, and our finding linking 3-HK to lysosomal acidification, we hypothesized that KAR antagonism could be promoting lysosomal acidification at least as part of its mechanism to promote bacterial clearance and host survival. We thus tested the impact of KAR antagonism and agonism on lysosomal acidification. Indeed, treatment of infected larvae with the KAR antagonist NS 3763 increased Lysotracker staining in infected macrophages *in vivo*, phenocopying exogenous 3-HK and confirming that KAR antagonism promotes lysosomal acidification in infected macrophages (Fig. 4B). Conversely, KAR agonism with the known agonist kainate had the opposite effect, decreasing Lysotracker staining in infected macrophages thereby phenocopying the defect in lysosomal acidification in Kmo inhibitor treated and knockdown animals (Fig. 4C).

Finally, as we have shown that KAR antagonism reverses the sensitivity to infection due to Kmo inhibition either chemically (1) or through genetic knockdown (Fig. 4A), we tested whether KAR antagonism reverses the defect in lysosomal acidification seen in macrophages in Kmo inhibitor treated and knockdown animals. The KAR antagonist NS 3763 indeed restored lysosomal acidification levels in both infected and uninfected Kmo inhibited macrophages, as measured using Lysotracker staining (Fig. 4D, Fig. S7D). Thus, 3-HK acts through KARs to promote lysosomal acidification and bacterial clearance in macrophages, revealing a novel mechanism by which a tryptophan metabolite acts through a neurotransmitter receptor as part of the innate immune response to bacterial infection.

## Discussion

The kynurenine pathway and specifically Kmo and its enzymatic product 3-hydroxy-kynurenine are required for defense against systemic *S.* Typhimurium infection in zebrafish larvae. When endogenous production of 3-HK is inhibited, macrophage microbicidal activity is impaired, resulting in greater bacterial numbers per macrophage and poor control of bacterial burden in the entire organism later in infection. 3-HK works via KARs to promote lysosomal acidification resulting in increased clearance of intracellular bacteria and thus improved host survival.

Loss of Kmo results in a global defect in lysosomal acidification in both infected and uninfected macrophages (Fig. 3D, Fig. S7A), which might suggest that lysosomal biogenesis itself is impaired following *kmo* knockdown. Indeed, the promoter regions of lysosomal genes found to be suppressed following *kmo* knockdown possess coordinated lysosomal expression and regulation (CLEAR) elements, which are recognized by the MiT/TFE family of transcription factors, including TFEB, which control lysosome biogenesis (24–26) (27). Interestingly, TFEB is regulated by external stimuli such as infection. For example, *Salmonella* has been previously shown to promote its own intracellular survival by perturbing lysosomal biogenesis through both direct inactivation as well as downregulation of TFEB (28). We observed a global suppression of lysosomal genes with CLEAR elements that are regulated by TFEB in our infected macrophage RNAseq dataset when *kmo* is knocked down, including vacuolar ATPase subunits *atp6ap1b*, *atp6v0b*, *atp6v0ca*; acidic hydrolases such as *ctsk*, *gaa2*, *aga*, *ctns*; and lysosomal membrane and trafficking proteins *laptm4a*, *mcoln1a*, and *ap1m1*. Hence, endogenous 3-HK may indirectly influence TFEB activation, which could account for why loss of Kmo activity impairs lysosomal acidification.

We had previously shown that neither treatment with the Kmo inhibitor Ro61-8048 or exogenous application of 3-HK or the KAR antagonist NS 3763 had any impact on *S.* Typhimurium infection in an array of cultured cell lines and primary cells typically used to model *Salmonella* infection, including macrophages (1). The lack of any phenotype in these reductionist models suggests that endogenous 3-HK promotion of lysosomal acidification through KARs may be a cell-extrinsic, systemic phenomenon not replicated in a dish. Hence, we propose that endogenous 3-HK antagonizes KAR channels in an as yet undefined cell type (*e.g.*, neurons, where KARs are most strongly expressed) to indirectly promote lysosomal acidification in macrophages in a systemic manner (Fig. 5). KAR activity had not been connected to infection outcomes before our previous work (1). Here, we expand upon the mechanism by which KAR antagonism improves survival to bacterial infection by linking KAR antagonism to increased lysosomal acidification in macrophages, which improves survival to systemic bacterial infection.

**Figure 5:**
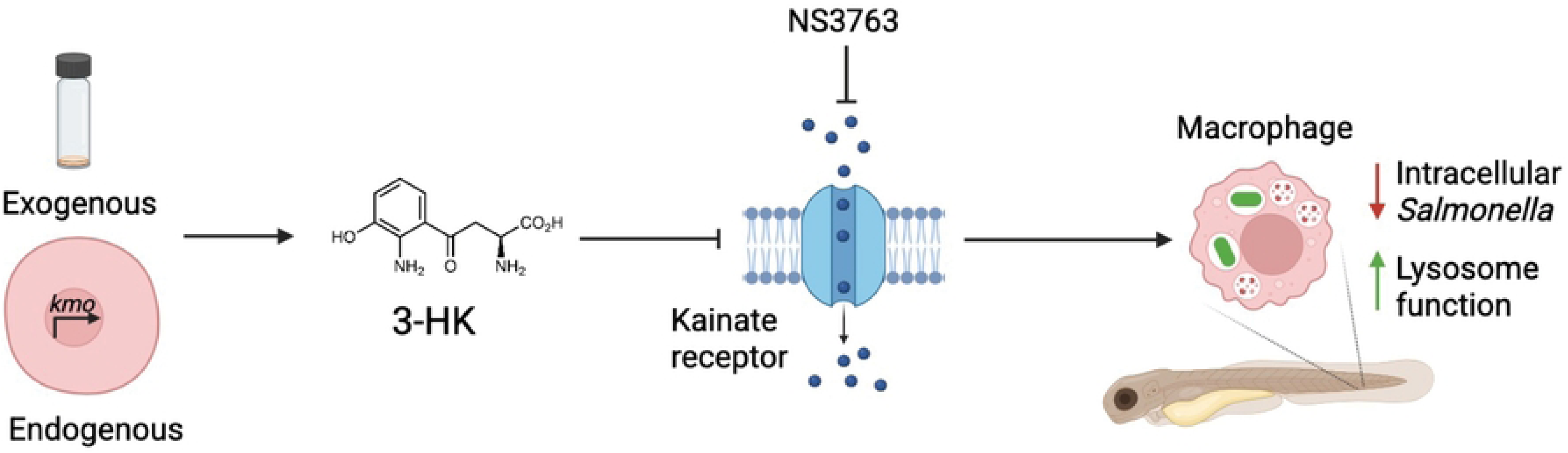
M**o**del **of the mechanism of action of 3-HK improving macrophage function during *in vivo* infection.** 3-HK, given exogenously or produced endogenously in the kynurenine pathway by Kmo, antagonizes kainate receptors to reduce the intracellular burden of *S*. Typhimurium in macrophages through enhanced lysosomal acidification. Schematic created with BioRender.com.

The kynurenine pathway has been implicated in the control of a number of pathogens including bacteria, viruses and parasites by a variety of mechanisms. For obligate intracellular tryptophan auxotrophs such as *Toxoplasma gondii* and some Chlamydial species, Ido-induced tryptophan deprivation is the primary mechanism by which the kynurenine pathway exerts its control over these pathogens as restoration of tryptophan prototrophy in these organisms relieves Ido1-mediated control (7, 29–31). For other organisms, the kynurenine pathway control of pathogen burden has been attributed to ‘toxic effects’ of downstream metabolites. For a few pathogens such as *L. monocytogenes* and *T. cruzi*, several downstream metabolites, including 3-HK, have direct effects on pathogen viability *in vitro* either resulting in bacterial cell killing (8) or ultrastructural changes that may inhibit intracellular pathogen replication (32). However, in these instances, the toxic effect (spectrum of activity) of these metabolites *in vitro* appear to be relatively restricted as these molecules do not have direct antimicrobial effects on other pathogens such as *S.* Typhimurium, *Pseudomonas aeruginosa*, *Listeria innocua* or *Streptococcus pyogenes* (1, 8). Here we reveal a novel mechanism of kynurenine pathway mediated control of systemic *S.* Typhimurium infection, with 3-HK activity at KARs promoting lysosomal acidification and control of bacterial burden in macrophages. This mechanism may be relevent to other bacterial and viral infections where loss of Kmo has been shown to be detrimental to survival (33, 34).

## Materials and Methods

### Bacterial strains

*Salmonella enterica* serovar Typhimurium SL1344 (gift from Ramnik Xavier, MGH) and *S*. Typhimurium-GFP (35) were utilized to infect zebrafish larvae. Strains were prepared by shaking bacterial inoculation overnight in LB. The overnight culture was then sub-cultured (1:500) for 3 hours at 37°C with shaking to mid-log phase. Bacteria were pelleted, washed 1X in PBS, and resuspended at 4X inoculum size in 1X PBS with 20% glycerol and frozen into single use aliquots at –80°C. Individual aliquots were thawed, diluted in PBS and 0.05% phenol red for infection.

### Zebrafish Lines

Wildtype zebrafish from the AB line and the transgenic line, Tg(*mpeg1.1:mCherry*)(gl22Tg) with mCherry-labeled macrophages under the Mpeg1.1 (macrophage expressed gene 1) promoter (19) were used in this study. Zebrafish were maintained in compliance with the guidelines and IACUC protocol approved by MGH’s Institutional Animal Care and Use Committee.

### Zebrafish infections

Zebrafish larvae (72 hpf) were inoculated intravenously by microinjection into the duct of Cuvier as previously described (1, 36–38). Infected larvae were washed 1X in embryo media (E3) then immersed in E3 containing DMSO vehicle control or compound in 96-well plates. Survival was measured at 24 hour intervals until 5 days post-infection. Bacterial inoculum was determined by plating the dose of injected bacteria onto agar plates and CFU was quantified following overnight incubation.

### Whole larvae bacterial enumeration

Individual infected larvae were isolated and then euthanized with ice bathing and excess tricaine for 5 minutes. Larvae were washed once with PBS then homogenized with shaking. To homogenize, larvae were placed in a microcentrifuge tube with a sterile 5 mm steal bead. Larvae were then shaken three rounds with each time consisting of 2 minutes at 40 Hz using a TissueLyser bead mill (Qiagen). Homogenized larvae were plated on streptomycin (25ug/mL) agar plates and CFU were determined by manual counting after overnight incubation.

### Small molecules

Kmo inhibitor Ro61-8048 from Tocris (99.3%). 3-hydroxy-D,L-kynurenine from BOC Sciences (>98%) or Biosynth Carbosynth (98%). Kainic acid from Tocris (>99.8%). NS 3763 prepared in house.

### Genetic knockdown

The translation-blocking *kmo* morpholino was purchased from Gene Tools and reconstituted to 1mM in sterile water (39). The morpholino was diluted to 100uM and 1-2 nL was microinjected into the one-to four-cell stage. CRISPR mutagenesis was adapted from previous work (40). Complexed guide RNAs (gRNA) were created by complexing equimolar amounts of Alt-R tracrRNA and designed kmo-specific Alt-R crRNAs (AA: GCCAGACGTACATTCCCCAT exon 7, AB: GATCTGATCGTAGGCTGTGA exon 7, AC: GGTTTGCTCTGTCCACGGAG exon 5). crRNA and tracrRNA was complexed in nuclease-free Duplex Buffer (IDT) and stored at –20 C until use. A week before use, gRNA was complexed with Alt-R Sp Cas9 Nuclease V3 (IDT) at an equimolar ratio in working buffer (20mM HEPES, 150mM KCl, pH 7.5) at 37 C for 5 minutes. Complexed RNPs were then pooled for each gene and 2 nL were injected into the 1-cell stage of wild-type AB zebrafish. CRISPR efficacy was assessed through next generation sequencing of PCR amplicons spanning CRISPR targets.

### Immune cell quantification

Macrophages were quantified by fluorescent microscopy and flow cytometry analysis. For microscopy, 3 dpf mpeg1:mCherry larvae from treated and untreated conditions were anesthetized with tricaine, immobilized in 1% low-melting-point agarose (Sigma) containing Tricaine in glass-bottom dishes (MatTek) and imaged on an epifluorescence microscope (Nikon 90i epifluorescence microscope). The entire larvae was imaged and macrophages were quantified using ImageJ (41). For flow cytometry analysis, larvae were dissociated at 4 HPI as previously described (42). Briefly, 5 pooled larvae were digested via mechanical contusion with trypsin and collagenase then assessed using flow cytometry (Cytek Aurora). Data was analyzed with Flowjo and Graphpad Prism.

### pHrodo staining

To assess the lysosome capability of Kmo-inhibitor treated larvae, pHrodo-labelled heat-killed *E.coli* particles (ThermoFisher) was injected into the bloodstream through the duct of Cuvier. At 4 HPI, 5 larvae were pooled for each biological replicate and dissociated as described above. Cell homogenates were filtered using a 40uM filter and analyzed using flow cytometry (NovoCyte 3000, Agilent) and Flowjo analysis.

### Lysotracker staining

Lysosome acidity was assessed using Lysotracker staining and analyzed with either flow cytometry or confocal microscopy. For flow cytometry analysis, Lysotracker staining of larvae was achieved on a modified protocol as previously described (43). Briefly, infected larvae were immobilized using Tricaine and 1 nL of Lysotracker Deep Red (Thermofisher) was injected into the Duct of Cuvier at 4 HPI. Infected larvae were sorted into groups of 5 larvae per biological replicate and staining was assessed following dissociation by flow cytometry (Cytek Aurora). Data was analyzed using Flowjo.

For confocal microscopy, larvae were infected then stained for 1 hour with 1 uM Lysotracker Deep Red (Thermofisher) in E3 in the dark at 3 HPI (44–46). At 4 HPI, after 1 hour of Lysotracker incubation, larvae were washed 3x in E3. Larvae were then immobilized using Tricaine and mounted after 10 minutes in low melting point agarose with 0.2 mg/mL Tricaine in 35 mm glass bottom dishes (MatTek) and imaged on Nikon AXR confocal microscope. Co-localization was assessed using Imaris.

### Transcriptomic analyses

Bulk RNA sequencing of macrophages was performed using an adapted Smart-seq2 protocol (47). For each biological replicate, 5 Tg(mpeg1:mCherry) larvae were dissociated as described above and single-positive mCherry+ macrophages or double positive mCherry+/GFP+ macrophages were isolated via FACS. cDNA libraries were prepared with Smart-Seq2 protocol in combination with Nextera XT Library preparation kit (Illumina Inc). Samples were multiplexed and pair end sequences were generated with NovaSeq SP sequencing system (Illumina) generating 5.5e10^7 reads per sample. Samples were demultiplexed and analyzed as before (1). Gene set enrichment analysis was performed with ClusterProfiler (v.4.0) with a FDR <0.05 using the Kyoto Encyclopedia of Genes and Genomes terms (48, 49).

### ROS production

Whole larvae ROS production was assessed as previously described using the permeant ROS indicator CM-H2DCFDA (Life Technologies) (50). Immediately following infection, larvae were placed in an opaque 96-well plate (Nunc 265301) in wells containing E3 or Ro61-8048 or 1mM Na-ascorbate as a positive antioxidant control. One hour after infection, CM-H2DFCDA was added to each well at a final concentration of 1ug/mL with 1% DMSO. Larvae were incubated with CM-H2DFCDA for 1 hour in the dark at 28C and then washed 1X with E3. Larvae were then monitored in the Tecan Spark 10M plate reader (Tecan) overnight at 28C with reads every hour at 484/529 excitation/emission. Wells without fish were used as a negative control.

### qRT-PCR

Gene expression was assessed through qRT-PCR. 8 larvae per sample were pooled at 16-19 HPI and euthanized with Tricaine and ice immersion for 5 minutes. Pooled larvae were homogenized in RLT buffer (Qiagen) using a 27 gauge needle and syringe. RNA was then isolated using RNeasy Min Elute Clean-up kit (Qiagen) according to manufacturer’s instructions with on-column DNA digestion. RNA concentration was assessed using Nanodrop and cDNA synthesis was performed with 2 ug RNA. mRNA expression levels were determined by quantitative real-time PCR using iQSYBR Green Supermix (BioRad). The following primers were used:

Kmo: ACTTGGACAGGACGTTCACA, TCAGAGCCTCGACTCCTATC

EF1A: ACCTACCCTCCTCTTGGTCG, GGAACGGTGTGATTGAGGGA

IL-1B: TAACCAGCTCTGAAATGATGGC, CCCTCATGCATGTCGCATCT

TNFa: TGCTTCACGCTCCATAAGACC, CAAGCCACCTGAAGAAAAGG

### Quantifying Phagocytosis and Intracellular Bacterial Load

To quantify phagocytosis of bacteria by macrophages, Tg(*mpeg1.1:mCherry*) larvae were infected with *S*. Typhimurium-GFP. At 4 HPI, larvae were dissociated using trypsin and collagenase as described above, with 5 larvae pooled per biological replicate. Homogenates were filtered using a 40uM filter, then the percentage of macrophages with bacteria was evaluated by flow cytometry (Cytek Aurora). To quantify intracellular bacterial load, Tg(*mpeg1.1:mCherry*) larvae were infected with *S*. Typhimurium-GFP and dissociated in groups of 5 larvae per biological replicate. Infected macrophages were then sorted using FACS (Aria 3) into a lysis buffer of PBS with 0.1% Triton X-100. Lysates were vortexed and bacterial CFU was enumerated by plating on LB-carbenicillin (100ug/mL) plates and counting resulting colonies after overnight incubation at 37C.

### Transmission Electron Microscopy

Tg(*mpeg1.1:mCherry*) larvae were infected with *S*. Typhimurium-GFP into the caudal vein. At 4 HPI larvae were screened for bacteria in the bloodstream through circulating GFP+ bacteria with fluorescent microscopy. Larvae with circulating bacteria were then euthanized on ice with Tricaine and fixed for electron microscopy. For transmission electron microscopy analysis, entire larvae were fixed overnight at 4C using 2.5% glutaraldehyde, 1.25% paraformaldehyde, and 0.03% picric acid in 0.1M sodium cacodylate buffer (pH 7.4). Larvae were then washed in 0.1M cacodylate buffer and postfixed with 1% osmium tetroxide and 1.5% potassium ferrocyanide for 1 hour, washed two times in water, once with maleate buffer, and then incubated for 1 hour in maleate buffer with 1% uranyl acetate. Larvae were washed twice in water and dehydrated using grades of ethanol. To embed larvae, larvae were incubated in propyleneoxide for 1 hour then infiltrated overnight in a 1:1 mixture of propylenoxide and TAAB (TAAB Laboratories Equipment, Ltd). The next day, the larvae were embedded in TAAB Epon and polymerized at 60 C for 48 hours. Ultrathin sagittal sections of the caudal hematopoietic tissue were cut using a Reichert Ultracut-S microtome. Sections were then stained with lead citrate and examined using a JEOL 1200EX transmission electron microscope. Images were recorded with an AMT 2k CCD camera.

## Acknowledgements

This work was funded by a generous gift from Anita and Josh Bekenstein (DTH) and a Fund for Medical Discovery Postdoctoral Fellowship from MGH (MPK).

## Supporting Information

**Supplemental 1: Validation of Kmo knockdown methods.** (**A**) *kmo* expression is significantly reduced in *kmo*-/-larvae compared to wild-type (Unpaired t-test. ****p<0.0001). **(B, C**) Inhibition of endogenous 3-HK production through Kmo depletion, using a morpholino to *kmo* (**B**), or Kmo inhibition, using Ro61-8048 (**C**), significantly sensitize larvae to infection, which can be rescued by exogenous addition of 3-HK. (**B**) *kmo* and control morphants were infected with *S.* Typhimurium (CFU= 228 +/− 32) and either treated with 100µM 3-HK or left untreated and monitored for survival (N= 14 larvae/ group; ****p<0.0001, Log rank test). (**C**) Larvae were infected with *S.* Typhimurium (CFU = 158 +/− 71) and treated with vehicle (DMSO), 50µM Ro61-8048, or 50µM Ro61-8048 and 100µM 3-HK and monitored for survival (N= 12 larvae/ group; *p<0.05, ***p<0.001, Log rank test). Data are representative of 3 experiments. (**D**) Kmo depletion impairs larval ability to control bacterial burden. Kmo and control morphant larvae were infected with *S.* Typhimurium (CFU= 1115 +/− 190) and bacteria were enumerated from individual larvae at indicated time points. Data are representative of 3 experiments (Unpaired t-test. **p<0.01).

**Supplemental 2: Kmo inhibition does not impair the inflammatory response to infection.** (**A**) Kmo inhibition with 50µM Ro61-8048 enhances ROS production over time in infected larvae. Larvae were infected with *S*. Typhimurium and ROS production was measured over time using the general oxidative stress indicator CM-H_2_DCFDA on a plate reader (Values are mean +/− SEM of 8 larvae/ group). (**B**) *il-1β* and *tnfα* expression is significantly increased by RT-qPCR at 19 HPI following infection and treatment with 50µM of the Kmo inhibitor Ro61-8048 compared to larvae treated with vehicle (DMSO) control. Data is representative of 2 experiments. (Values are mean +/− SEM. Unpaired t-test ***p<0.001, **p<0.01).

**Supplemental 3: Kmo inhibition or depletion does not alter macrophage numbers or phagocytosis.** (**A-C**) Kmo depletion or inhibition does not alter macrophage numbers. (**A**) Tg(*mpeg1.1:mCherry*) Kmo morphant and control larvae were infected with *S.* Typhimurium or left uninfected. mCherry+ macrophage numbers were quantified by microscopy 17 HPI (**A**) or by flow cytometry 4 HPI (**B**). (**C**) Tg(*mpeg1.1:mCherry*) larvae were infected with *S.* Typhimurium, treated with 50µM Ro61-8048 or vehicle (DMSO) control and mCherry+ macrophage numbers were quantified by flow cytometry 4 HPI (**C**). (Values are mean +/− SEM. Unpaired t-test, ns, non-significant). (**D-F**) Kmo depletion or inhibition does not impair macrophage phagocytic engulfment. (**D,E**) Tg(*mpeg1.1:mCherry*) Kmo and control morphant larvae were injected with *S.* Typhimurium-GFP (**D**) or GFP+ latex beads (**E**) and the number of mCherry+/GFP+ macrophages were quantified by flow cytometry 4 HPI (Unpaired t-test, ns, non-significant; *, p<0.05). (**F)** Tg(*mpeg1.1:mCherry*) larvae were injected with GFP+ latex beads and treated with 50µM Ro61-8048 or vehicle (DMSO) control and the number of mCherry+/GFP+ macrophages quantified by flow cytometry 4 HPI (Values are mean +/− SEM; Unpaired t-test; ns, not significant). Data is representative of 3 replicates.

**Supplemental 4: Gating strategy for flow cytometry and FACS of infected macrophages.** Representative flow cytometry dot plots of single cells with gates for uninfected macrophages (DAPI^-^, mCherry^+^, GFP^-^) and infected macrophages (DAPI^-^, mCherry^+^, GFP^+^).

**Supplemental 5: Intracellular *S*. Typhimurium in double-membrane vacuoles.** (A) Representative images showing the presence of double membranes indicated with black arrows surrounding engulfed bacteria in *kmo* morphant and control morphant phagocytes at 4 HPI from transmission electron microscopy images of the caudal hematopoietic tissue.

**Supplemental 6: Infected macrophages show transcriptional signatures of infection, immune, and inflammatory pathways compared to naïve macrophages using bulk RNAseq.** Comparison of transcriptional changes measured using bulk RNAseq between infected and naïve macrophages isolated from control morphant larvae 4 HPI using gene set enrichment analysis. GO terms shows expected increases in infection, immune, and inflammatory pathways (**A**). (**B**) Fold-changes in individual genes of the defense response GO term in infected versus uninfected macrophages are displayed.

**Supplemental 7: Kmo inhibition or depletion impairs lysosome acidification in both exposed and infected macrophages.** (**A**) Kmo inhibition with Ro61-8048 decreased maximum Lysotracker fluorescence in macrophages, regardless of infection status, which was restored upon treatment with exogenous 3-HK. Tg(*mpeg1.1:mCherry*) larvae were infected with *S.* Typhimurium-GFP, treated with vehicle (DMSO) control, 50µM Ro61-8048, or 50µM Ro61-8048 and 100µM 3-HK. Lysosomal acidification was quantified by measuring Lysotracker staining at 4 HPI in macrophages using confocal microscopy (Values are individual macrophages from 8-13 biological replicates; One-way ANOVA followed by Tukey’s multiple comparisons. *p<0.05, ****p<0.0001.) (**B, C**) Kmo inhibition (B) or depletion (**C**) decreases bright pHrodo staining in macrophages. Heat-killed green pHrodo-*E.coli* particles were injected into Tg(*mpeg1.1:mCherry*) larvae treated with 50µM Ro61-8048 or vehicle (DMSO) control (**B**) or into Tg(*mpeg1.1:mCherry*) Kmo or control morphant larvae (**C**) and the number of pHrodo+/mCherry+ macrophages were quantified by flow cytometry at 4 HPI (**B,C**, each value represents 5 pooled larvae; unpaired t-test, **p<0.01, ***p<0.001;). (D) Decreased Lysotracker staining in macrophages following Kmo inhibition with Ro61-8048 is restored with the KAR antagonist NS 3763. Tg(*mpeg1.1:mCherry*) larvae were infected with *S.* Typhimurium-GFP, treated with vehicle (DMSO) control, 50µM Ro61-8048 or 50µM Ro61-8048 and 75µM of the KAR antagonist NS 3763. Lysosomal acidification was quantified by measuring Lysotracker staining in macrophages using confocal microscopy (Values are individual macrophages from 8-13 biological replicates; One-way ANOVA followed by Tukey’s multiple comparisons, ****p<0.0001.)

**Supplemental 8: 3-HK requires lysosomal acidification to improve survival to infection.** Treatment with Bafilomycin A1, a vATPase inhibitor that also inhibits autophagy, abrogates the ability of 3-HK to rescue larvae from infection. Larvae were infected with *S.* Typhimurium (238 +/− 88 CFU), treated with 125nM Bafilomycin A1, 100µM 3-HK, or DMSO control and monitored over time for survival. (N= 15 larvae/ group; ****p<0.0001, Log rank test).

## References

1. Parada-Kusz M, Clatworthy AE, Goering ER, Blackwood SM, Shigeta JY, Mashin E, et al. 3-Hydroxykynurenine targets kainate receptors to promote defense against infection. Nat Chem Biol. 2024.

2. Platten M, Nollen EAA, Rohrig UF, Fallarino F, Opitz CA. Tryptophan metabolism as a common therapeutic target in cancer, neurodegeneration and beyond. Nat Rev Drug Discov. 2019;18(5):379–401.

3. Ozaki Y, Edelstein MP, Duch DS. The actions of interferon and antiinflammatory agents of induction of indoleamine 2,3-dioxygenase in human peripheral blood monocytes. Biochem Biophys Res Commun. 1987;144(3):1147–53.

4. Pfefferkorn ER, Rebhun S, Eckel M. Characterization of an indoleamine 2,3-dioxygenase induced by gamma-interferon in cultured human fibroblasts. J Interferon Res. 1986;6(3):267–79.

5. Opitz CA, Litzenburger UM, Lutz C, Lanz TV, Tritschler I, Koppel A, et al. Toll-like receptor engagement enhances the immunosuppressive properties of human bone marrow-derived mesenchymal stem cells by inducing indoleamine-2,3-dioxygenase-1 via interferon-beta and protein kinase R. Stem Cells. 2009;27(4):909–19.

6. Wang Y, Liu H, McKenzie G, Witting PK, Stasch JP, Hahn M, et al. Kynurenine is an endothelium-derived relaxing factor produced during inflammation. Nat Med. 2010;16(3):279–85.

7. McClarty G, Caldwell HD, Nelson DE. Chlamydial interferon gamma immune evasion influences infection tropism. Curr Opin Microbiol. 2007;10(1):47–51.

8. Nino-Castro A, Abdullah Z, Popov A, Thabet Y, Beyer M, Knolle P, et al. The IDO1-induced kynurenines play a major role in the antimicrobial effect of human myeloid cells against Listeria monocytogenes. Innate Immun. 2014;20(4):401–11.

9. Knubel CP, Martinez FF, Fretes RE, Diaz Lujan C, Theumer MG, Cervi L, et al. Indoleamine 2,3-dioxigenase (IDO) is critical for host resistance against Trypanosoma cruzi. FASEB J. 2010;24(8):2689–701.

10. Chandrakar P, Parmar N, Descoteaux A, Kar S. Differential Induction of SOCS Isoforms by Leishmania donovani Impairs Macrophage-T Cell Cross-Talk and Host Defense. J Immunol. 2020;204(3):596–610.

11. Fields PI, Swanson RV, Haidaris CG, Heffron F. Mutants of Salmonella typhimurium that cannot survive within the macrophage are avirulent. Proceedings of the National Academy of Sciences of the United States of America. 1986;83(14):5189–93.

12. Leung KY, Finlay BB. Intracellular replication is essential for the virulence of Salmonella typhimurium. Proc Natl Acad Sci U S A. 1991;88(24):11470–4.

13. Trede NS, Langenau DM, Traver D, Look AT, Zon LI. The use of zebrafish to understand immunity. Immunity. 2004;20(4):367–79.

14. van der Sar AM, Musters RJ, van Eeden FJ, Appelmelk BJ, Vandenbroucke-Grauls CM, Bitter W. Zebrafish embryos as a model host for the real time analysis of Salmonella typhimurium infections. Cell Microbiol. 2003;5(9):601–11.

15. Stockhammer OW, Zakrzewska A, Hegedus Z, Spaink HP, Meijer AH. Transcriptome profiling and functional analyses of the zebrafish embryonic innate immune response to Salmonella infection. J Immunol. 2009;182(9):5641–53.

16. Hall CJ, Flores MV, Oehlers SH, Sanderson LE, Lam EY, Crosier KE, et al. Infection-responsive expansion of the hematopoietic stem and progenitor cell compartment in zebrafish is dependent upon inducible nitric oxide. Cell Stem Cell. 2012;10(2):198–209.

17. Gogoi M, Shreenivas MM, Chakravortty D. Hoodwinking the Big-Eater to Prosper: The Salmonella-Macrophage Paradigm. J Innate Immun. 2019;11(3):289–99.

18. Masud S, Prajsnar TK, Torraca V, Lamers GEM, Benning M, Van Der Vaart M, et al. Macrophages target Salmonella by Lc3-associated phagocytosis in a systemic infection model. Autophagy. 2019;15(5):796–812.

19. Ellett F, Pase L, Hayman JW, Andrianopoulos A, Lieschke GJ. mpeg1 promoter transgenes direct macrophage-lineage expression in zebrafish. Blood. 2011;117(4):e49–56.

20. Murayama E, Kissa K, Zapata A, Mordelet E, Briolat V, Lin HF, et al. Tracing hematopoietic precursor migration to successive hematopoietic organs during zebrafish development. Immunity. 2006;25(6):963–75.

21. Ordas A, Hegedus Z, Henkel CV, Stockhammer OW, Butler D, Jansen HJ, et al. Deep sequencing of the innate immune transcriptomic response of zebrafish embryos to Salmonella infection. Fish Shellfish Immunol. 2011;31(5):716–24.

22. Mauvezin C, Neufeld TP. Bafilomycin A1 disrupts autophagic flux by inhibiting both V-ATPase-dependent acidification and Ca-P60A/SERCA-dependent autophagosome-lysosome fusion. Autophagy. 2015;11(8):1437–8.

23. Lerma J, Marques JM. Kainate receptors in health and disease. Neuron. 2013;80(2):292–311.

24. Sardiello M, Palmieri M, di Ronza A, Medina DL, Valenza M, Gennarino VA, et al. A gene network regulating lysosomal biogenesis and function. Science. 2009;325(5939):473–7.

25. Palmieri M, Impey S, Kang H, di Ronza A, Pelz C, Sardiello M, et al. Characterization of the CLEAR network reveals an integrated control of cellular clearance pathways. Hum Mol Genet. 2011;20(19):3852–66.

26. La Spina M, Contreras PS, Rissone A, Meena NK, Jeong E, Martina JA. MiT/TFE Family of Transcription Factors: An Evolutionary Perspective. Front Cell Dev Biol. 2020;8:609683.

27. Tan A, Prasad R, Lee C, Jho EH. Past, present, and future perspectives of transcription factor EB (TFEB): mechanisms of regulation and association with disease. Cell Death Differ. 2022;29(8):1433–49.

28. Rao S, Xu T, Xia Y, Zhang H. Salmonella and S. aureus Escape From the Clearance of Macrophages via Controlling TFEB. Front Microbiol. 2020;11:573844.

29. Nelson DE, Virok DP, Wood H, Roshick C, Johnson RM, Whitmire WM, et al. Chlamydial IFN-gamma immune evasion is linked to host infection tropism. Proc Natl Acad Sci U S A. 2005;102(30):10658–63.

30. Pfefferkorn ER. Interferon gamma blocks the growth of Toxoplasma gondii in human fibroblasts by inducing the host cells to degrade tryptophan. Proceedings of the National Academy of Sciences of the United States of America. 1984;81(3):908–12.

31. Sibley LD, Messina M, Niesman IR. Stable DNA transformation in the obligate intracellular parasite Toxoplasma gondii by complementation of tryptophan auxotrophy. Proc Natl Acad Sci U S A. 1994;91(12):5508–12.

32. Knubel CP, Insfran C, Martinez FF, Diaz Lujan C, Fretes RE, Theumer MG, et al. 3-Hydroxykynurenine, a Tryptophan Metabolite Generated during the Infection, Is Active Against Trypanosoma cruzi. ACS Med Chem Lett. 2017;8(7):757–61.

33. Zhao J, Chen J, Wang C, Liu Y, Li M, Li Y, et al. Kynurenine-3-monooxygenase (KMO) broadly inhibits viral infections via triggering NMDAR/Ca2+ influx and CaMKII/ IRF3-mediated IFN-beta production. PLoS Pathog. 2022;18(3):e1010366.

34. Hoshi M, Kubo H, Ando T, Tashita C, Nakamoto K, Yamamoto Y, et al. 3-Hydroxykynurenine Regulates Lipopolysaccharide-Stimulated IL-6 Production and Protects against Endotoxic Shock in Mice. Immunohorizons. 2021;5(6):523–34.

35. Avraham R, Haseley N, Brown D, Penaranda C, Jijon HB, Trombetta JJ, et al. Pathogen Cell-to-Cell Variability Drives Heterogeneity in Host Immune Responses. Cell. 2015;162(6):1309–21.

36. Clatworthy AE, Lee JS, Leibman M, Kostun Z, Davidson AJ, Hung DT. Pseudomonas aeruginosa infection of zebrafish involves both host and pathogen determinants. Infect Immun. 2009;77(4):1293–303.

37. Cui C, Benard EL, Kanwal Z, Stockhammer OW, van der Vaart M, Zakrzewska A, et al. Infectious disease modeling and innate immune function in zebrafish embryos. Methods Cell Biol. 2011;105:273–308.

38. Benard EL, van der Sar AM, Ellett F, Lieschke GJ, Spaink HP, Meijer AH. Infection of zebrafish embryos with intracellular bacterial pathogens. J Vis Exp. 2012(61).

39. Korstanje R, Deutsch K, Bolanos-Palmieri P, Hanke N, Schroder P, Staggs L, et al. Loss of Kynurenine 3-Mono-oxygenase Causes Proteinuria. J Am Soc Nephrol. 2016;27(11):3271–7.

40. Kroll F, Powell GT, Ghosh M, Gestri G, Antinucci P, Hearn TJ, et al. A simple and effective F0 knockout method for rapid screening of behaviour and other complex phenotypes. Elife. 2021;10.

41. Schindelin J, Arganda-Carreras I, Frise E, Kaynig V, Longair M, Pietzsch T, et al. Fiji: an open-source platform for biological-image analysis. Nat Methods. 2012;9(7):676-82.

42. Gallardo VE, Liang J, Behra M, Elkahloun A, Villablanca EJ, Russo V, et al. Molecular dissection of the migrating posterior lateral line primordium during early development in zebrafish. BMC Dev Biol. 2010;10:120.

43. Fan J, Hale VL, Lelieveld LT, Whitworth LJ, Busch-Nentwich EM, Troll M, et al. Gaucher disease protects against tuberculosis. Proc Natl Acad Sci U S A. 2023;120(7):e2217673120.

44. He C, Klionsky DJ. Analyzing autophagy in zebrafish. Autophagy. 2010;6(5):642–4.

45. Peri F, Nusslein-Volhard C. Live imaging of neuronal degradation by microglia reveals a role for v0-ATPase a1 in phagosomal fusion in vivo. Cell. 2008;133(5):916–27.

46. Lopez-Cuevas P, Xu C, Severn CE, Oates TCL, Cross SJ, Toye AM, et al. Macrophage Reprogramming with Anti-miR223-Loaded Artificial Protocells Enhances In Vivo Cancer Therapeutic Potential. Adv Sci (Weinh). 2022;9(35):e2202717.

47. Picelli S, Faridani OR, Bjorklund AK, Winberg G, Sagasser S, Sandberg R. Full-length RNA-seq from single cells using Smart-seq2. Nat Protoc. 2014;9(1):171–81.

48. Wu T, Hu E, Xu S, Chen M, Guo P, Dai Z, et al. clusterProfiler 4.0: A universal enrichment tool for interpreting omics data. Innovation (Camb). 2021;2(3):100141.

49. Yu G, Wang LG, Han Y, He QY. clusterProfiler: an R package for comparing biological themes among gene clusters. OMICS. 2012;16(5):284–7.

50. Roca FJ, Ramakrishnan L. TNF dually mediates resistance and susceptibility to mycobacteria via mitochondrial reactive oxygen species. Cell. 2013;153(3):521–34.

